# Molecular characterization of effector protein SAP54 in *Bellis Virescence Phytoplasma* (16SrIII-J)

**DOI:** 10.1101/411140

**Authors:** Franco D Fernández, Humberto J Debat, Luis R Conci

## Abstract

Phytoplasmas are wall-less bacteria, with a parasitic life style responsible for numerous plant diseases worldwide. The genomic landscape of phytoplasmas has been scarcely studied. Nevertheless, since the introduction of next generation sequencing technologies, genome wide studies of these pathogens are flourishing and a handful of phytoplasmas genomes are available in public databases. In South America, phytoplasmas from 16SrIII group (X-disease) are the most widely distributed, and only a draft genome from a phytoplasmas 16SrIII-J infected periwinkle from Chile has been generated (Phytoplasma Vc33). Here, in grafting experiments we characterized the phenotypic signatures of an Argentinian daisy derived isolate of a 16SrIII-J phytoplasma (*Bellis Virescence* Phytoplasma) infecting periwinkle. Moreover, we applied a pipeline for genome wide annotation of the Vc33 genome and identified the effector protein SAP54. We then employed the obtained data to amplify, clone, sequence and characterize a SAP54 orthologue protein of *Bellis Virescence* Phytoplasma. Structural and phylogenetic analyses suggested that the identified SAP54 is highly conserved, and that its co-divergence among phytoplasma is not directly consistent with the evolutionary trajectories derived from rRNA analyses. The results gathered here could provide the basis for reverse genetics experiments using 16SrIII-J SAP54 proteins to assess their eventual influence in pathogenesis.

## 1. Introduction

Phytoplasmas are wall-less bacteria, with parasitic life style responsible for numerous plant diseases worldwide (Hogenhout et al. 2008; Lee et al. 2000; Maejima et al. 2014a). More than 1,000 plant species have been reported to be affected by these pathogens including crops, ornamental plants and trees (Bertaccini et al. 2014). Phytoplasmas are obligate parasites of plant phloem and of insect vectors. Within plants they have a habitat restricted to the sieve elements of the phloem where they grow and multiply (Christensen et al. 2004). Despite numerous efforts, it has been challenging to obtain stable *in vitro* phytoplasma cultures (Zhao et al. 2015), which limited the study of this kind of pathogen. Hence, basic forward or reverse genetics strategies to characterize their gene repertoire have been unattainable. In this scenario, most studies dealing with phytoplasmas have relied in genomic annotation approaches to infer and understand basic aspects of pathogenicity of these bacteria. While five genomes of phytoplasmas have been reported based on traditional Sanger technologies (Oshima et al. 2004; Bai et al. 2006; Tran-Nguyen et al. 2008, Kube et al. 2008, Andersen et al. 2013), since the implementation of next generation sequencing (NGS) methods, a rapid increase of available genome data has scaled. Since NGS, one additional complete genome has been depicted (Orlovskis et al. 2017), and other 16 have been partially (draft) sequenced (Chung et al. 2013; Mitrović et al. 2014; Saccardo et al. 2012, Fischer et al. 2016; Pacifico et al. 2015; Zamorano and Fiore 2016; Sparks et al. 2018). Analyzes of these genomes have shown, despite their small size, the phytoplasma genome contains a substantial number of genes in multiple copies (Bai et al. 2006; Davis et al. 2003; Jomantiene and Davis 2006; Oshima et al. 2004). Most of these genes are organized in clusters dubbed potential mobile units (PMUs) (Bai et al. 2006) or sequence-variable mosaics (SVMs) (Jomantiene et al. 2007; Oshima et al. 2004). In these regions, genes have been identified for membrane-bound proteins and most of the effector protein genes, both associated with virulence factors (Hogenhout and Bos 2011; Toruño et al. 2010).

It has been hypothesized that effector proteins of phytoplasmas induce morphological and physiological changes in host plants, which could increase pathogen fitness during infection (Sugio et al. 2011). For instance, these phytoplasmas secreted proteins could produce phyllody, a symptoms that causes conversion of floral organs into green leaf-like organs, which may prolong the life span of their hosts, since as biotrophic pathogens phytoplasmas require a host to live (Sugio et al. 2011). Recently, it was shown that a family of highly conserved phytoplasma virulence factors, designated as a phyllody-inducing gene family (Phyllogen, SAP54 homologues), induce phyllody and other floral malformations in *Arabidopsis thaliana* (MacLean et al. 2011; Maejima, et al. 2014b; Yang et al. 2015). In addition, a recent report has shown that Phyllogen can induce phyllody in several species of angiosperms (Kitazawa et al. 2017), so these proteins have been proposed to play a central role as key factors involved in the generalized virulence of phytoplasmas.

In South America phytoplasmas from 16SrIII group (X-disease) are the most widely distributed and with the broader host range. Within this group, phytoplasmas belonging to subgroup 16SrIII-J were associated to diverse diseases in Argentina, Chile and Brazil (Fernández et al. 2017; Fugita et al. 2017; Montano et al. 2011). Plant species affected by this subgroup are diverse and include garlic (*Allium sativum*), chayote (*Sechium edule*), sunflower (*Helianthus annuus*), cauliflower (*Brassica oleracea*), eggplant (*Solanum melongena*), strawberry (*Fragaria* × *ananassa*), daisy (*Bellis perennis*), woolflowers (*Celosia spp*.), periwinkle (*Catharanthus roseus*), parrot fruit (*Aegiphila verticillata*), ironweed (*Vernonia brasiliana*), lettuce (*Lactuca sativa*), Swiss chard (*Beta vulgaris*) and cassava (*Manihot esculenta*) (Montano et al. 2011; Pérez-López et al. 2016; Quiroga et al. 2017; Fernández et al. 2018). Despite the growing importance of this particular subgroup of phytoplasmas, only one draft genome has been obtained so far (Zamorano and Fiore 2016). Moreover, there are no reports describing effector proteins in phytoplasmas from subgroup 16SrIII-J present in Argentina. In the present work, we used the available genomic information in order to detect and characterize for the first time the SAP54 protein sequence in a phytoplasma isolate of subgroup 16SrIII-J in Argentina.

## 2. Materials and methods

### 2.1 Plant material, graft transmission and symptomatology evaluation

*Bellis virescence* Phytoplasma (*BellVir*) is a phytoplasma belonging to subgroup 16SrIII-J previously described in association to virescence and phyllody on daisy (*Bellis perennis*) (Galdeano et al., 2013). This phytoplasma was maintained and propagated in periwinkle (*Catharathus roseus*) from daisy (*Bellis perennis*) infected plants by grafting. In this work we used *BellVir* phytoplasma as representative strain of subgroup 16SrIII-J. In order to evaluate the symptomatology developed by *BellVir* infection, 10 periwinkle plants (5 of white flowers, 5 of pink flowers) (2-3 months old) agamically obtained by rooting cuttings, were grafted with this phytoplasma. The periwinkle plants came from two mother plants (1 white flower, 1 pink flower) so they constituted an isogenic line. The grafted plants were maintained in humid chambers for 15 days. After this period, the plants were kept under controlled conditions of temperature (25°C) and humidity (RH 60-80%) in greenhouses. Evolution of symptomatology was evaluated by visual inspection, focusing attention on phyllody and virescence symptoms. Phytoplasmas detection was evaluated by PCR using the X-disease group (16SrIII) specific primers P1/Xint (Smart et al. 1996), which amplify a fragment ∼1.6kb of 16S rRNA operon. DNA used as template was isolated from petioles and midribs of grafted periwinkles (symptomatics, 3-5 months after graft) and healthy periwinkles were ground in liquid nitrogen and DNA purified according to Doyle & Doyle (1990). PCR reactions were performed in solutions containing 100 ng of DNA, 0.4 mM of each primer, 200 mM of each dNTP, 1 U of GoTaq^®^ DNA polymerase, 1X polymerase buffer (Promega, USA) and sterile water to a final volume of 40 μl. PCR amplifications were evaluated trough electrophoresis in agarose gel stained with GelRed^®^ (Biotium, USA).

### 2.2 Detection and characterization of effector protein SAP54 homolog in 16SrIII-J reference genome

So far, only one draft genome has been described for the subgroup 16SrIII-J (Zamorano and Fiore, 2016). The complete sequences (29 contigs) were retrieved from Genbank (accession number: LLKK00000000.1, Bioproject PRJNA293833) and CDSs were annotated using the Rapid Annotation using Subsystem Technology (RAST) server (http://rast.nmpdr.org). Among the total predicted proteins, those with presence of signal peptide (predicted using Signal IP 4.1 as implemented in http://www.cbs.dtu.dk/services/SignalP/ using the sensitive D-cutoff values) and without transmembrane domains outside the signal peptide (predicted with the TMHMM Server v. 2.0 available at http://www.cbs.dtu.dk/services/TMHMM/) were identified. Proteins which passed this filter, and thus considered as putative secreted proteins, were analyzed in order to identify SVM (Sequence-variable mosaic) protein signals (Pfam id: 12113) using the Conserved Domains Database search tool (www.ncbi.nlm.nih.gov/Structure/cdd/wrpsb.cgi, expect value = 0.01, CDD v3.16 database). Also, a BLASTp search targeting proteins with SVM-signal (consensus sequence: MFKLKNQLLIINIFLFIFLGLFLITNNNQVMAM, E-value ≤ 1e-05) against a local dataset (encompassing all the CDSs annotated by RAST) was performed in Geneious R.10 (Biomatters, USA). The final set of proteins (Signal Peptide (+); Transmebrana domains outside SP (-) and SVM-signal (+)) were analyzed by reciprocal BLASTp searches (E-value ≤ 1e-05) against Aster yellows witches’-broom phytoplasma AYWB proteins (taxid:322098) for identification of SAPs homologs (Bai et al. 2006).

### 2.3 PCR Amplification and sequencing of the predicted SAP54 gene from BellVir (16SrIII-J) phytoplasma

Amplification of DNA sequence of *BellVir* SAP54-LIKE gene was acceded by conventional PCR. A DNA contig harboring a SAP54-LIKE gene was identified and *de novo* primer design was made using Primer 3 (http://primer3.ut.ee/). PCR reactions were made using DNA from phytoplasma positive and healthy periwinkle plants as a template as previously described in 2.1 (conditions of amplification are described in Supplementary materials). PCR amplification products were purified using GFX-columns (GE Healthcare, USA) and cloned in pGEM-T easy system (Promega, USA) according to manufacturer. The cloned fragments were bi-directionally Sanger sequenced (Macrogen, Korea). A final consensus sequence (3X coverage) was assembled using Geneiuos R.10 software and deposited in Genbank database (NCBI accession number MH756633).

### 2.4 Molecular characterization of SAP54-LIKE gene in BellVir phytoplasma

Consensus sequence of SAP54-LIKE gene was analyzed using Geneious R.10. The SAP54 homologous protein sequence was obtained and structural analysis was conducted as previously described in 2.2 (signal peptide, Transmembrane domains, SVM-signal). BLASTp (nr-database) analysis was performed in order to identify putative SAP54-homologues in NCBI database. Also, phylogenetic relationships were inferred using the SAP54 reference sequences (protein) and 16S rDNA corresponding sequence. Multiple alignments were conducted employing MAFFT (L-INS-i algorithm, BLOSUM62 scoring matrix) and MUSCLE (clustering method: UPGMB, max interations: 8) for SAP54 and 16rDNA sequences respectively. Phylogenetic threes were constructed using the Maximum-Likelihood method (JTT model; gamma distributed, NNI heuristic method) in MEGA6 software (Tamura et al. 2013).

## 3. Results and Discussion

### 3.1 BellVir phytoplasma infection in periwinkle

After 3 months post-graft, plants began to show the first symptoms of virescence. In both white flowers and pink flowers, this symptom was characterized by the presence of flowers with green spots (Figure 1.b, e). After the appearance of this symptom the plants began to develop the characteristic phyllody symptom (Figure 1.c, f). The symptomatology of filodia remained constant until the plants ceased the production of flowers and began to show additional symptoms of chlorosis, shortening of internodes and decrease in leaf size. Total grafted plants (10/10) presented the described symptomatology, and by PCR the presence of phytoplasmas was verified in all the plants as well. No amplification was observed in healthy control plant (Figure 1.a, d).

**Figure 1:**
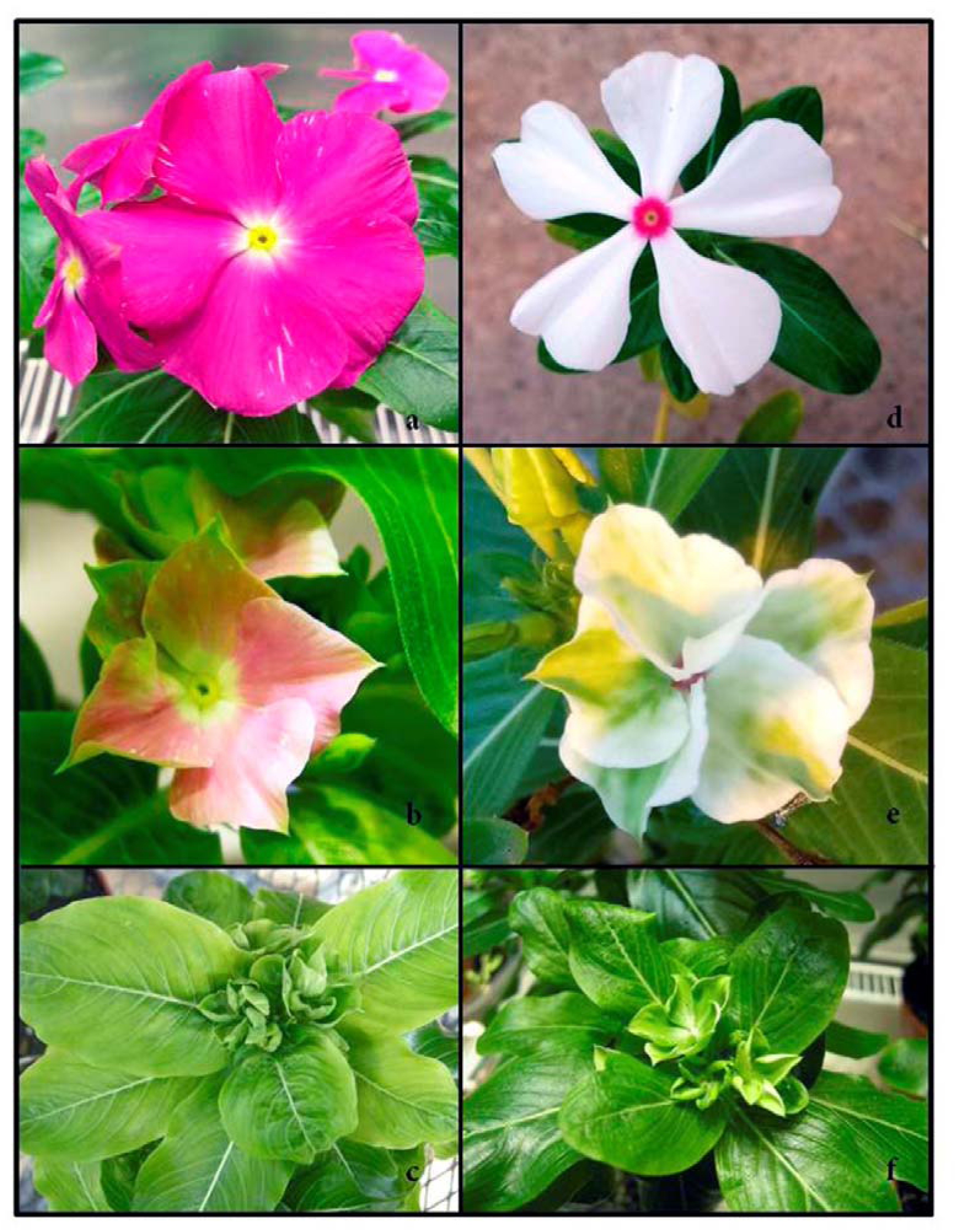
Symptomatology associated to infection of *BellVir* phytoplasma in grafted periwinkles. 1. a, d: healthy flowers; 1.b,e: symptoms of virescence and 1.c,f: symptoms of phyllody.

### 3.2 Identification of SAP54 homologue in 16SrIII-J phytoplasma genome

During the annotation process of the draft genome of the periwinkle associated 16SrIII-J phytoplasma, Zamorano and Fiore (2016) detected 11 CDSs containing the SEC translocase complex signal peptide, but the authors stated that they did not find any evidence of SAP11, SAP54, PHYL, or TENGU like genes. As these genes and their products appear to be fundamentals in the pathogenic process of phytoplasma infection, we embarked in an exhaustive exploration of their sequence dataset in order to assess the presence of signatures associated with these important genes. Using the RAST platform a total of 634 CDSs and 30 RNAs sequences were identified in the 16SrIII-J phytoplasma draft genome (Table S1, Supplementary material). The predicted protein sequences of these 634 CDSs were analyzed by Signal IP (gram +, sensitive) and peptide signal was detected in 54 sequences. Putative Secreted Proteins (PSP=no transmembrane domains after signal-peptide) were identified in 37 out 54 SP-proteins. SVM-signal was identified by CDD search (NCBI) or by sequence homology (BLASTp-search against local database) in seven CDSs (Table 1). Identification of putative SAP homologues proteins was acceded by BLASTp analysis of these seven putative effector proteins against the Aster Yellows Witches Broom phytoplasma reference genome (NC_007716.1) (Bai et al. 2006). All in all, our pipeline succeeded in the identification, with high confidence, of a putative SAP54 (AYWB_224) homologue (protein ID: fig|33926.32.peg.561, Table 2) in the 16SrIII-J genome. The encoded protein had 114 aa in length (13.410 kDa) and showed a 56% aa identity (E-value: 3,00e-24, coverage 85%) with the SAP54 reference sequence (Table 2, ABC65341.1).

**Table 1:**
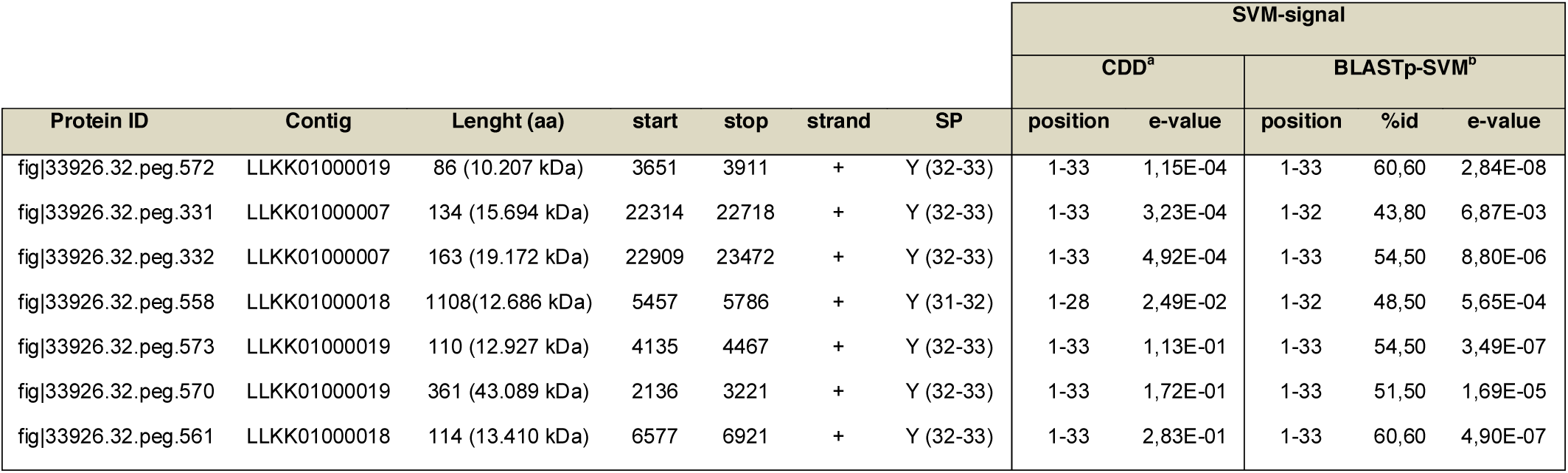
Identification Putative secreted proteins (PSP) of Vc33 Phytoplasma (16SrIII-J) CDSs. Protein IDs were assigned by RAST (See supplementary table 1). a: position and e-value of SVM-signal (pfam12113) using the Conserve Domains Database search tool; b: position, identity percentages and e-values of PSP against SVM consensus domain using BLASTp (local database).

| Protein ID | Contig | Lenght (aa) | start | stop | strand | SP | SVM-signal |  |  |  |  |
| --- | --- | --- | --- | --- | --- | --- | --- | --- | --- | --- | --- |
|  |  |  |  |  |  |  | CDD<sup>a</sup> |  | BLASTp-SVM<sup>b</sup> |  |  |
|  |  |  |  |  |  |  | position | e-value | position | %id | e-value |
| fig\|33926.32.peg.572 | LLKK01000019 | 86 (10.207 kDa) | 3651 | 3911 | + | Y (32-33) | 1-33 | 1,15E-04 | 1-33 | 60,60 | 2,84E-08 |
| fig\|33926.32.peg.331 | LLKK01000007 | 134 (15.694 kDa) | 22314 | 22718 | + | Y (32-33) | 1-33 | 3,23E-04 | 1-32 | 43,80 | 6,87E-03 |
| fig\|33926.32.peg.332 | LLKK01000007 | 163 (19.172 kDa) | 22909 | 23472 | + | Y (32-33) | 1-33 | 4,92E-04 | 1-33 | 54,50 | 8,80E-06 |
| fig\|33926.32.peg.558 | LLKK01000018 | 1108(12.686 kDa) | 5457 | 5786 | + | Y (31-32) | 1-28 | 2,49E-02 | 1-32 | 48,50 | 5,65E-04 |
| fig\|33926.32.peg.573 | LLKK01000019 | 110 (12.927 kDa) | 4135 | 4467 | + | Y (32-33) | 1-33 | 1,13E-01 | 1-33 | 54,50 | 3,49E-07 |
| fig\|33926.32.peg.570 | LLKK01000019 | 361 (43.089 kDa) | 2136 | 3221 | + | Y (32-33) | 1-33 | 1,72E-01 | 1-33 | 51,50 | 1,69E-05 |
| fig\|33926.32.peg.561 | LLKK01000018 | 114 (13.410 kDa) | 6577 | 6921 | + | Y (32-33) | 1-33 | 2,83E-01 | 1-33 | 60,60 | 4,90E-07 |

**Table 2:** Identification of SAP (Bai et al., 2006) protein homologues in Putative Secreted Proteins of Vc33 Phytoplasma (16SrIII-J).* BLASTp analysis was conducted in Geneious R.10 software against the genome of AYWB (NC_007716.1) in local database. Putative SAP54 is marked in bold.

| Protein ID | BLASTp-AYWB* |  |  |  |  | Locus_tag=Putative SAP |
| --- | --- | --- | --- | --- | --- | --- |
|  | Identity | Coverage | E-value | Accession | Product |  |
| fig\|33926.32.peg.572 | 60% | 83% | 3,00E-24 | ABC65485.1 | conserved hypothetical protein | AYWB_368=SAP67 |
| fig\|33926.32.peg.331 | 56% | 98% | 1,00E-44 | ABC65149.1 | conserved hypothetical protein | AYWB_032=SAP05 |
| fig\|33926.32.peg.332 | 72% | 99% | 8,00E-63 | ABC65190.1 | conserved hypothetical protein | AYWB_073=SAP19 |
| fig\|33926.32.peg.558 | 59% | 100% | 1,00E-23 | ABC65150.1 | conserved hypothetical protein | AYWB_033=SAP06 |
| fig\|33926.32.peg.573 | 44% | 66% | 1,20E-09 | ABC65354.1 | conserved hypothetical protein | AYWB_237=SAP41 |
| fig\|33926.32.peg.570 | 58% | 92% | 5,00E-67 | ABC65456.1 | conserved hypothetical protein | AYWB_339=SAP49 |
| **fig\|33926.32.peg.561** | **56%** | **85%** | **3,00E-24** | ABC65341.1 | **conserved hypothetical protein** | **AYWB_224=SAP54** |

### 3.3 Sequencing and analysis of SAP54 gen of BellVir phytoplasma

Once the sequence of the putative SAP54 protein in the 16SrIII-J reference genome was predicted, primers were designed for the amplification by PCR to confirm our *in silico* findings. A new set of primers (SAP54Fw1/SAP54Rv1) (Supplementary material) was designed in order to amplify a fragment of ∼615 bp containing the putative SAP54 gene (Figure 2.a). Using these primers, products of ∼615 bp were visualized in gel electrophoresis of PCR amplifications with DNA from all periwinkle plants (10/10) infected with *BellVir* phytoplasma, and no PCR products were evident in DNA obtained from uninfected periwinkle plants. Amplifications products from two infected plants were bi-directionally Sanger sequenced and compared sharing a 100% identity. A final consensus sequence was deposited in Genbank with MH756633 accession number. The assembled sequence (615 bp) contains a single ORF (+1frame, position 41-394) of 354 nt which encodes a protein of 117 aa. The sequenced *BellVir* protein shared an 83.3% aa identity with the 114 aa SAP54 protein predicted in the draft genome of periwinkle associated 16SrIII-J phytoplasma (Zamorano and Fiore 2016). This protein presented a signal peptide (SP, position 32-33) and no transmembrane domain after the SP. The SVM-signal (pfam12113) was also confirmed (e-value= 5.89e-06) from position 1 to 33 (MFRSKNQFKIIHLCLIAFIGLLFIFNNHQLMAM). A comparison with the SAP54 reference sequence (AYWB) showed an aa identity value of 51% and a conservation of the typical SAP54 structure (Figure 2.a-b). A multiple alignment using SAP54 homologues available from genbank, showed a consistent deletion of six aa (position 47-52) in *BellVir*, Vc33 phytoplasma (both from subgroup 16SrIII-J) and also from ‘*Ca*. Phytoplasma solani’ (subgroup 16SrXII-B). In addition, these three SAP54 homologues presented at the C-region a three aa insertion with a conserved NN(I/L) motif (Figure 2.b). These sequences (Vc33 and ‘*Ca*. Phytoplasma solani’) were also the most similar in terms of identity (Table 3). The C-terminal end (residues 106 to 129) was the most conserved throughout all the compared sequences. Interestingly, this region is significantly associated with coiled-coils structural motifs (Figure 2.a-b).

**Table 3:** List of sequences used in Phylogenetic analyses. *: aminoacidic identity values against putative SAP54 of BellVir phytoplasma (MH756633).

| Candidatus Phytoplasma | Strain | 16Sr-subgroup | Protein ID | %id* | Reference |
| --- | --- | --- | --- | --- | --- |
| *Ca. Phytoplasma pruni* | Vc33 | 16SrIII-J | LLKK010000018 | 83.33 | Zamorano and Fiore 2016 |
| *Ca. Phytoplasma solani* | 284-09 | 16SrXII-B | CCP88716 | 71.79 | Mitrovic et al., 2014 |
| *Ca. Phytoplasma pruni* | JR1 | 16SrIII-A | WP_01791948.1 | 52.85 | Saccardo et al., 2012 |
| *Ca. Phytoplasma asteris* | OY-M | 16SrI-B | BAD04134 | 48.80 | Oshima et al., 2004 |
| *Ca. Phytoplasma asteris* | OY-V | 16SrI-B | GAK74327 | 48.80 | Kakizawa et al., 2014 |
| *Ca. Phytoplasma asteris* | CY-P | 16SrI-B | WP_034172429 | 46.40 | Pacifico et al., 2015 |
| *Ca. Phytoplasma asteris* | AYWB | 16SrI-A | ABC65341 | 48.00 | Bai et al., 2006 |
| *Ca. Phytoplasma trifolii* | 16SrVI-A | 16SrVI-A | BAQ08267 | 48.00 | Maejima et al., 2014 |
| - | PnWB-NTU2011 | 16SrII-A | EMR14798 | 40.32 | Chung et al., 2013 |
| - | EPWB-NCHU2014 | 16SrII-A | WP_004994552.1 | 40.32 | Chang et al. 2015 |
| *Ca. Phytoplasma Phoenicium* | ChiP | 16SrIX-B | PQP79243 | 46.40 | Quaglino et al. 2015 |

**Figure 2:**
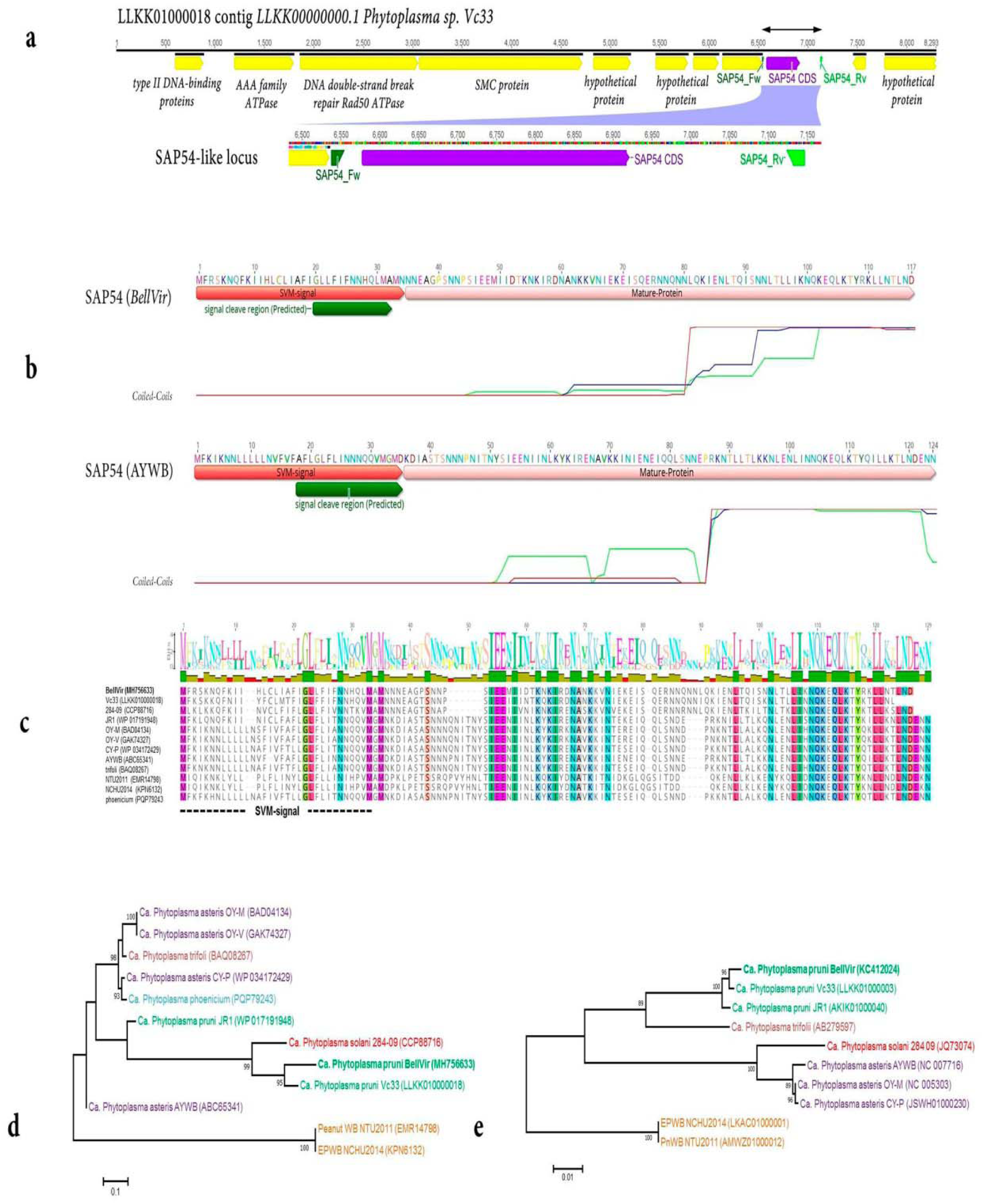
Analyses of SAP54 protein sequence of *BellVir* phytoplasma. **a:** Genomic context of SAP54 CDS in LLKK01000018 contig, primer binding sites are marked in green. **b**: comparison of SAP54 structure between *BellVir* phytoplasma (MH756633) and AYWB phytoplasma (ABC65341); probabilities of coiled-coil regions (MTK matrices) are presented under sequence structure (windows 14-21-28: green, blue and red respectively). **c**: multiple alignment of SAP54 homologues proteins, identical residues are marked in colors, SVM in dashed line. **d** and **e**: Phylogenetic trees of SAP54 and 16S rDNA sequences using ML method. The different ‘*Candidatus* Phytoplasma species’ are differentiated with colors. The GenBank accession number for each taxon is given in parentheses. The numbers on the branches are bootstrap (confidence) values (1,000 replicates). The scale bar represents the number of amino acid substitutions per site.

Phylogenetic analyses were conducted from both SAP54 protein sequences and 16S rDNA sequences (Figures 2.c-d). The 16S rDNA is the most widely used gene for evaluation of phylogenetic relationships and taxonomic assignment in phytoplasmas (Lee et al., 1998; 2000). For SAP54 protein sequences, a maximum likelihood tree showed that *BellVir* and Vc33 were grouped in the same clade with the ‘*Ca*. Phytoplasma solani’ (Figure 2.d). Interestingly, the general topology of this tree does not consistently correspond to that of the 16S gene, since the groupings generated do not share the same clustered taxonomic groups (Figure 2.d). Given the intrinsic nature of this virulent associated proteins and their influence in the pathogenic outcome of plant-pathogen interactions, it is tempting to suggest that the inconsistences in the evolutionary clustering of rRNA and SAP54 trees might be associated with selective pressure and active genomic drift of these specific proteins. Further studies, in the context of additional data, should assess genome based analyses of evolutionary trajectories of these pathogens.

## 4. Conclusions

In this work we demonstrated by grafting tests on periwinkle plants, that infection by *BellVir* (16SrIII-J) phytoplasma causes severe symptoms of virescence and phyllody. Previous studied suggest that such symptomatology could be triggered by a phyllody-inducing gene family (Phyllogen, SAP54 effector protein) (MacLean et al. 2011; Maejimaet al. 2014; Yang et al. 2015) in several species of angiosperms (Kitazawa et al. 2017). Among the diversity of phytoplasmas composing the 16SrIII-J clade, some strains have been also reported to induce virescence-phyllody symptoms in sunflower (Guzmán et al. 2014), squash (Galdeano et al. 2013) and in periwinkle (Pérez-López et al. 2016). These outcomes could be explained by the presence of SAP54 homologues proteins or alternative pathways. However, experimental data to confirm this hypothesis remains elusive. Zamorano and Fiore (2016) generated the first draft genome of a phytoplasma from subgroup 16SrIII-J. An exhaustive exploration of their sequence dataset allowed us to identify with high confidence a putative SAP54 (AYWB_224) homologue. We then employed this data to amplify, clone, sequence and characterize a SAP54 orthologous protein of *Bellis Virescence* Phytoplasma. The sequenced *BellVir* SAP54 protein (MH756633) conserved the typical structure of SAP54 (phyllogen) reference protein (Figure 2.b). The *BellVir* SAP54 protein (117aa-13.69 kDa) shared an 83.3% aa identity with the predicted SAP54 of Vc33 draft genome and 51% with the SAP54 reference sequence (ABC65341). Comparative analyses between SAP54 homologues showed that the C-terminal end is the most highly conserved region throughout all the assessed sequences. Interestingly, this region is significantly associated with coiled-coils structural motifs, which could provide new clues about the mechanisms of action of this effector. Phylogenetic analyses suggested that the identified SAP54 is highly conserved, and that its co-divergence among phytoplasma is not directly consistent with the evolutionary history derived from rRNA analyses. In the present work, the identification and molecular characterization of a SAP54 protein homologous in a phytoplasma belonging to subgroup 16SrIII-J was performed for the first time. The information generated here will be instrumental for further investigation of the eventual mechanisms of pathogenicity associated with this effector and thereby contribute to the overall knowledge of the pathosystem.

## BellVir sequences

~~~
>SAP54-LIKE_BellVir_DNA
ATGTTTCGATCAAAAAACCAATTTAAAATAATTCATCTTTGTTTAATCGCTTTTATAGGATTATTATTTATTTTTAATAATCATCAATTAATGGCGATGAATAATAATGAAGCTGGCCCAAGCAATAATCCATCAATTGAAGAAATGATTATTGATACAAAAAATAAAATTCGCGATAATGCAAATAAAAAAGTTAATATAGAAAAAGAAATATCACAAGAAAGAAATAATCAAAATAATCTTCAAAAAATTGAAAATCTTACTCAAATATCAAATAATT TAACATTATTAATTAAAAATCAAAAAGAACAACTAAAAACCTATAGAAAACTTTTAAATACTTTAAATGATTAA
>SA54-LIKE_BEllVir_aa
MFRSKNQFKIIHLCLIAFIGLLFIFNNHQLMAMNNNEAGPSNNPSIEEMIIDTKNKIRDNANKKVNIEKEISQERNNQNNLQKIENLTQISNNLTLLIKNQKEQLKTYRKLLNTLND
~~~

